# Fine-tuning Protein Embeddings for Generalizable Annotation Propagation

**DOI:** 10.1101/2023.06.22.546084

**Authors:** Andrew M. Dickson, Mohammad R. K. Mofrad

**Author notes:** The authors declare no competing interests.

## Abstract

A central goal of bioinformatics research is to understand proteins on a functional level, typically by extrapolating from experimental results with the protein sequence information. One strategy is to assume that proteins with similar sequences will also share function. This has the benefit of being interpretable; it gives a very clear idea of why a protein might have a particular function by comparing with the most similar reference example. However, direct machine learning classifiers now outperform pure sequence similarity methods in raw prediction ability. A hybrid method is to use pre-trained language models to create protein embeddings, and then indirectly predict protein function using their relative similarity. We find that fine-tuning an auxiliary objective on protein function indirectly improves these hybrid methods, to the point that they are in some cases better than direct classifiers. Our empirical results demonstrate that interpretable protein comparison models can be developed using fine-tuning techniques, without cost, or even with some benefit, to overall performance. K-nearest neighbors (KNN) embedding-based models also offer free generalization to previously unknown classes, while continuing to outperform only pre-trained models, further demonstrating the potential of fine-tuned embeddings outside of direct classification.

**Significance Statement:** For the purposes of biological understanding, or engineering, it is particularly important that protein annotation is done through interpretable means, such as comparison to existing proteins with known properties. Our work builds upon previous efforts to do so by embedding proteins into high-dimensional vectors with pre-trained protein language models. We demonstrate that further training of these models on protein function classification drastically improves their performance, while allowing for exactly the same similarity based classifications on both known and novel potential functions.

---

Despite decades of research, protein function is still only sparsely understood, with a huge percentage of proteins uncharacterized, or incompletely understood. However, the sequence information on amino acids making up proteins is typically known with high accuracy, meaning that any effective method for predicting functions from sequence would fill an enormous gap in biological research.

Datasets such as the Gene Ontology describe thousands of the possible functions for proteins in a comprehensive hierarchy, which works as a rigorous target for machine learning prediction (1). Proteins may be annotated with one or more GO terms independently, making functional annotation a multi-class, multi-label problem setting. GO annotation is characterized by an extremely high number of classes, but also by relative rarity of most classes. While the most frequent annotations are attached to most of the protein dataset, the vast majority occur only for a handful of samples. Out of the top 800 terms used for our experimental dataset, half of the terms occur in less than 0.2% of datapoints.

Heavy effort is put into predicting GO annotations for proteins, with trends towards fine-tuned language models as the strongest modern classifiers. However, the most popular methods for GO annotation in practice are based on sequence similarity, like BLAST (2, 3). Despite slightly lower accuracy, they have the advantage of explaining their predictions with examples of similar proteins with known function.

A simple way to reproduce this functionality with deep learning models is to make annotation predictions indirectly, by embedding proteins in some high-dimensional latent space and then classifying them with K-nearest neighbors using Euclidean distance as a similarity metric, an approach with precedents in text search, and protein homology prediction (4, 5). Here, we investigate the utility of this approach, in comparison with BLAST and with standard deep learning models, and demonstrate that it is highly effective under typical fine-tuning procedures (6). It joins a growing number of uses for self-supervised language models in modern proteomics (7–10).

For this problem setting, we derive an annotation dataset from existing databases. To limit the effect of missing annotations, we constrain our study to the set of human-reviewed proteins in SwissProt (11) and only train or evaluate on proteins with at least one annotation. Additionally, because annotation performance is dominated by performance on the most common gene ontology terms, we limit training to gene ontology terms which occur over 50 times in our database. The final dataset is generated with the gobench.org web application based on the Gene Ontology Annotation dataset (12), with similar methodology to the CAFA competition (13). In total, it contains 112,000 proteins, with an average of ten annotations for each protein.

## Results

### Fine-Tuning Attains Highest Performance for GO Annotation

As a reference baseline, we reproduce a subset of existing work in deep learning, fine-tuning pretrained models, and annotation through sequence comparison. We first use a modified version of BLAST based on prior work from DeepGOPlus (14) as a strong baseline. In practice, we find that it’s significantly stronger than typical BLAST approaches because it estimates protein annotation probabilities with a weighted average of similar proteins, instead of a direct comparison to the most similar protein.

We train convolutional models based on prior work in deep protein annotation (14, 15), and extensively hyperparameter tune them to maximize their strength as a baseline. We use the BERT-BFG model pre-trained by RostLab for fine-tuning(16), following a typical training process. The BERT model is converted to a classifier by condensing its final, internal sequence embeddings into a single vector, and passing them through a task-specific, linear classifier head. With the exception of the linear head, all model weights in the annotation model are transferred from the BERT model pre-trained to predict masked sequences. All model weights are fine-tuned for annotation prediction on a slanted triangular learning schedule (17).

We also use the pre-trained model, and a protein embedding given by the average of residue embeddings, to make annotations based on K-nearest neighbors. This follows prior work in annotating based only on language model embedding similarity (18).

Following the example of CAFA (13), all models are evaluated with precision-recall and MIS-RUS curves. Performance is summarized in the form of F-max and S-min metrics which give the F1 and S scores when model logit predictions are thresholded with the optimum value. Both are described in detail in an early CAFA paper (19). Because missing data is a significant problem for Gene Ontology datasets (20, 21), we take the extra step of calculating an ‘F-Estimate’ score, which is an estimate proportional to the true F1 score with no false negative labels (22). Encouragingly, relative F-estimate scores match F-max scores in nearly all cases, indicating that our testing dataset is a good proxy for real world performance.

Model results show that BLAST is extremely competitive, which matches previous results for Gene Ontology prediction datasets (14).

Our major overall takeaway is that the only classifier model that directly surpasses a tweaked BLAST is a fine-tuned language model with a classifier head. This improves over BLAST for both F-max and F-estimate scores, and is roughly the same for S-min scores, which put higher emphasis on rare GO classes (23). Although similarity search with pre-trained embeddings is highly desirable for interpretable results, without further training it seems that it’s initially weaker than most direct deep learning approaches.

### Fine-tuned KNN Models Outperform Direct Classifiers

Classification models are trained to output logit values for each Gene Ontology class in our dataset, so they can’t be used directly for similarity based comparisons. However, we find that internal model representations, especially the relatively small embedding directly before a classification head, work as effective representations for K-Nearest Neighbor classifiers. We adopt K-Nearest Neighbors to the multi-label setting by running it independently for each Gene Ontology class. The probability of a GO annotation is then estimated as the average number of neighbors associated to the annotation themselves. We find that performance is robust to the value of K, and fix a value of 10 for all experiments.

Convolutional models optimized for 128 or 2048 dimensions prior to the classifier layer achieve similar performance, both as classifiers and embedding models for KNN. With both representation sizes, KNN annotation has slightly lower performance than direct classification with model logits.

Conversely, for fine-tuned language models, KNNs based on the fine-tuned embedding are actually even more effective than the classifiers themselves, making them a more interpretable classification model with no drawbacks to performance. Investigation of model performance on individual GO classes shows that relying on a KNN improves performance significantly,

### Fine-tuned Embeddings Generalize to Unseen Classes

Fine-tuned BERT embeddings are stronger than those from the pre-trained version for propagating classifications with K-nearest neighbors. However, it is plausible that a fine-tuned model will have lost information critical to overall protein understanding in the process of being optimized for our specific target GO classes. We evaluate the significance of this by generating a train/test split not only across proteins, but also across GO classes, so that we can evaluate embedding for KNN performance on classes never previously used to train our model. We then use model embeddings, and when applicable model classifier predictions, to separately evaluate the F-max, S-min, and F-score performance metrics for GO classes included and excluded from training.

As we expect, the relative performances of each model is exactly the same for GO classes included in training data. Fine-tuned BERT remains the strongest model, with a KNN based on its internal embeddings still outperforming the base classification head.

For out-of-distribution classes, it’s no longer possible to evaluate the classification head, so all referenced models are based on either K-nearest neighbors or BLAST. The finetuned model continue to be much stronger than exclusively pre-trained embeddings, even for classes not included in training. The gap in F1 score is not directly interpretable, because one set of GO classes may be more difficult to classify than the other, but it is clear that fine-tuning continues to have a significant effect, without overfitting to include information only relevant to specific classes in training. It appears that fine-tuning successfully restricts models embeddings to a functionally relevant subspace, without significantly excluding details unrelated to known functions included in training data. However, without the benefit of training on known classes, direct sequence comparison with BLAST once again outperforms pure deep learning models.

### Model Performance on Out-of-Distribution Proteins

Within our testing dataset, most proteins have a handful of high significance (e-value less than 0.001) BLAST hits against the training dataset, which we consider to be strongly related (24). Prior models are evaluated on the restricted set of proteins with zero high significance matches against training data. This regime artificially lowers BLAST performance to zero.

Here, we find that pre-trained embeddings are now slightly stronger than most other models as evaluated by both F-max and S-min. As expected, all models trained on GO data suffer significantly, but fine-tuned models retain their overall advantage.

## Discussion

### Significance

In this work, we investigate the properties of finetuned language models for protein annotation, and we take the novel step of evaluating the impact of fine-tuning on representations for nearest neighborhood classification.

**Fig. 1.**
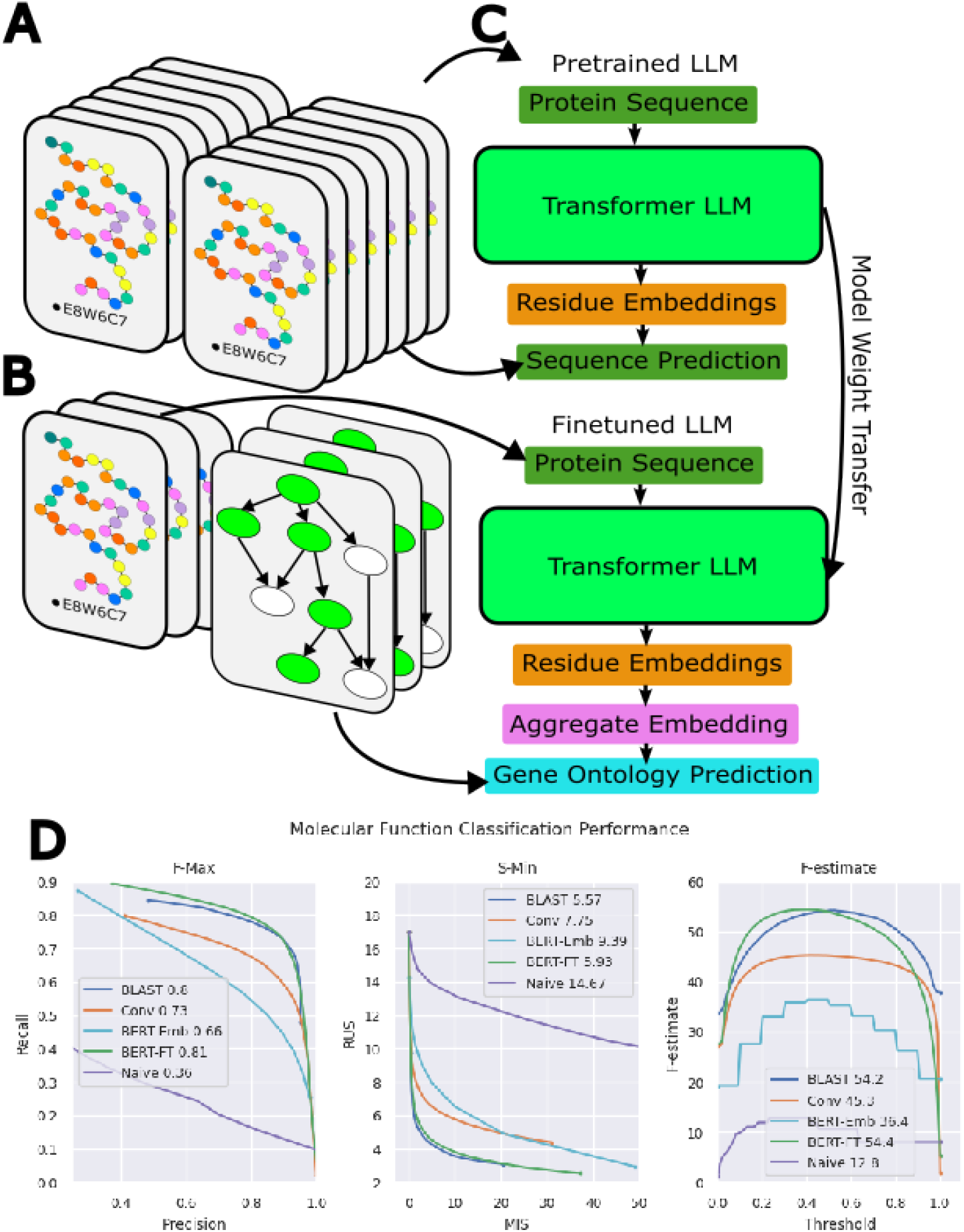
Initial benchmarking models. (A) Dataset structure for self-supervised language models. Pre-trained LLM’s are built to predict randomly masked residues of known protein sequences, with no auxilary data. Because of the simplicity of the dataset, it is much larger (3.83 hundred billion tokens) (16). (B) Dataset structure for fine-tuning on protein annotations. Dataset is built from protein sequences paired with a set of Gene Ontology annotations, organized hierarchically. The fine-tuning dataset is relatively small, with 112,000 proteins. (C) Deep-learning model architectures used for evaluation. The pre-trained LLM predicts protein sequences with the transformer architecture by predicting an embedding vector for each protein residue and putting each residue through a classifier head. Weights and architecture from the pre-trained model are transferred to an annotation model for fine-tuning. Here, the model is modified to average the residue embeddings and add them to the BERT CLS token embedding to form a sequence level representation, which is then fed into a classifier layer. Finally, a baseline convolutional model applies a larger set of increasingly large convolutions to the input protein sequence, and concatenates the results into a single, large, embedding vector. This is shrunk by several MLP layers to a bottleneck size of 2048 for our baseline model, and passed into a classifier. (D) Performance curves for each of our initial classifier models. Curves for naive class prediction based only on prior probabilities of GO classes are included, but because their performance is universally poor, they are excluded from further figures. Precision-recall and F-estimate curves show fine-tuned models as strictly stronger than alternatives, including BLAST, while MIS-RUS curves show BLAST as slightly stronger in some cases.

**Fig. 2.**
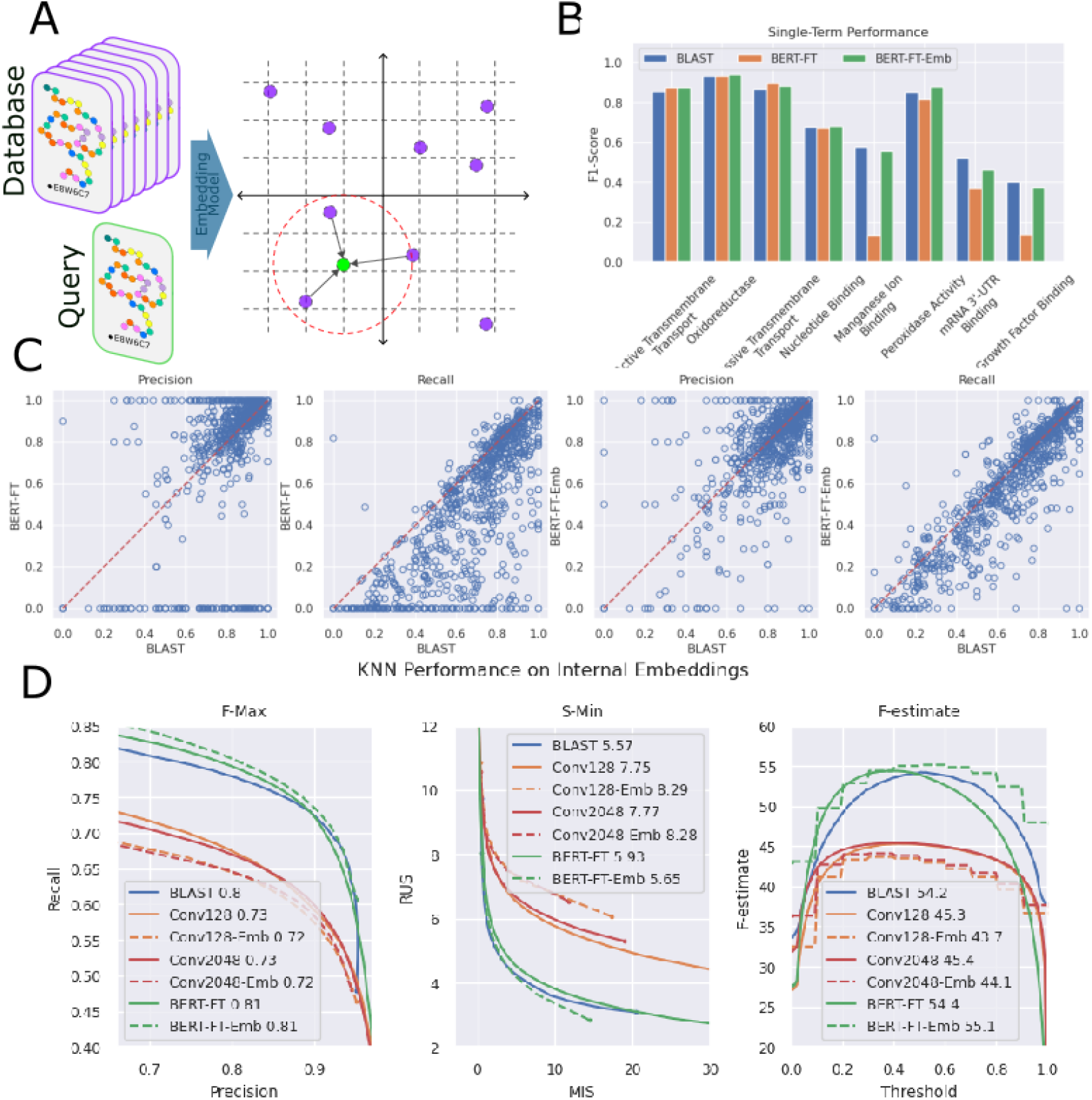
(A) Deep K-Nearest Neighbors scheme. Proteins in training data are embedded using some embedding model, either a large language model or a convolutional model in our results. Proteins in validation or testing dataset are then embedded by the same model, so that their closest neighbors from the training database can be queried. Annotation probability is then calculated as the average frequency of GO classes within nearest neighborhood. (B) Model F1 scores on selected specific GO classes, at p=0.5 threshold. (C) Scatterplots of precision or recall for each of 865 GO classes included in dataset. X-axis gives precision or recall for baseline BLAST model, while Y-axis shows corresponding value on same GO class for a fine-tuned or embedding model. Points below red line have higher precision or recall when annotated by BLAST. (D) Performance curves for classifiers relative to KNN for their internal embeddings. BLAST and pre-trained language model curves included for reference. Classifier models indicated with solid lines, while KNN on model internal embeddings indicated with dashed lines. Note that for fine-tuned embeddings, KNN curve is actually strictly higher than the trained classifier.

**Fig. 3.**
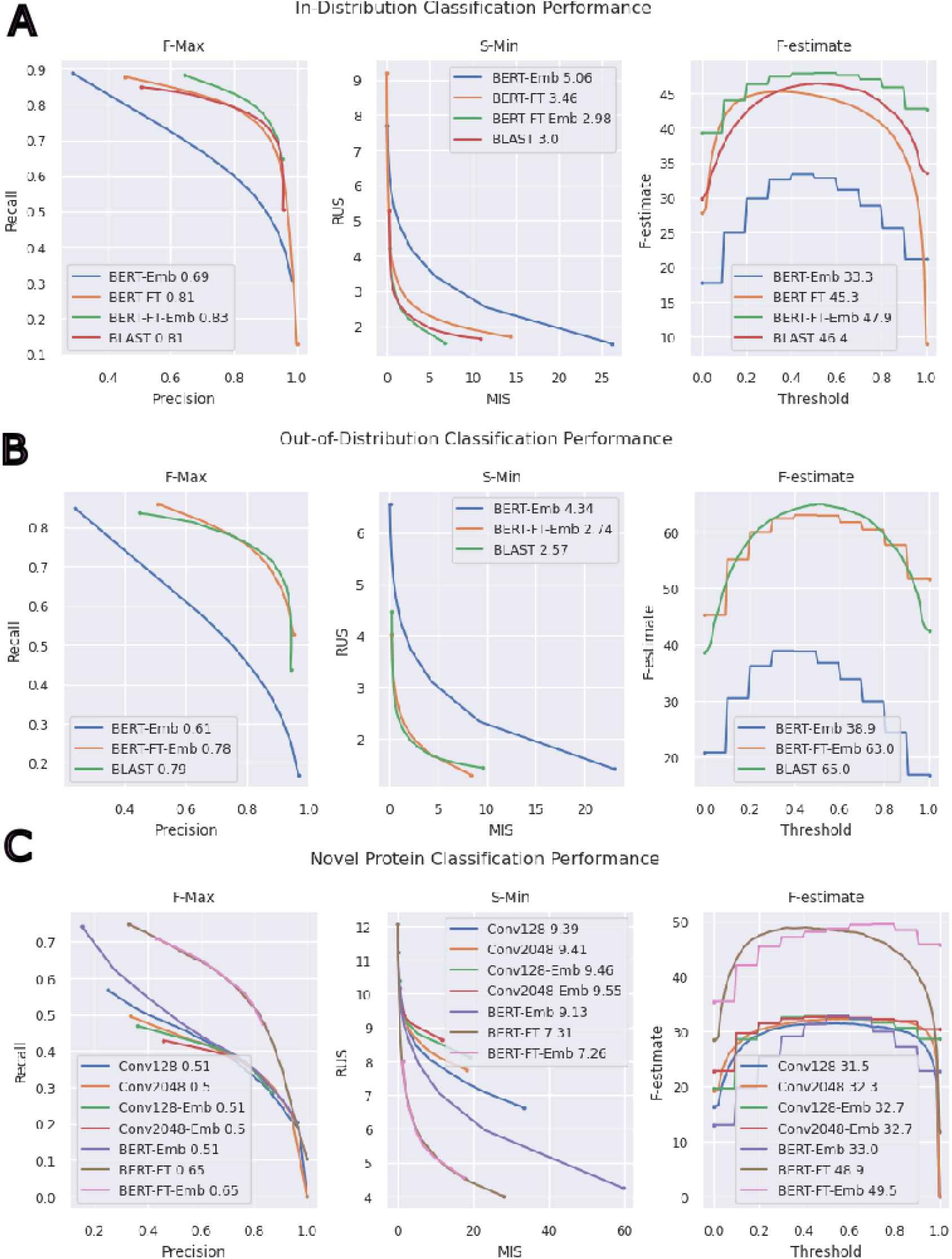
(A) Performance of selected models on 50% subset of GO terms used for training. (B) Performance of query based models on 50% subset of GO terms excluded from training. (C) Performance of all models on sequences with no significant BLAST matches in training data.

**Fig. 4.**
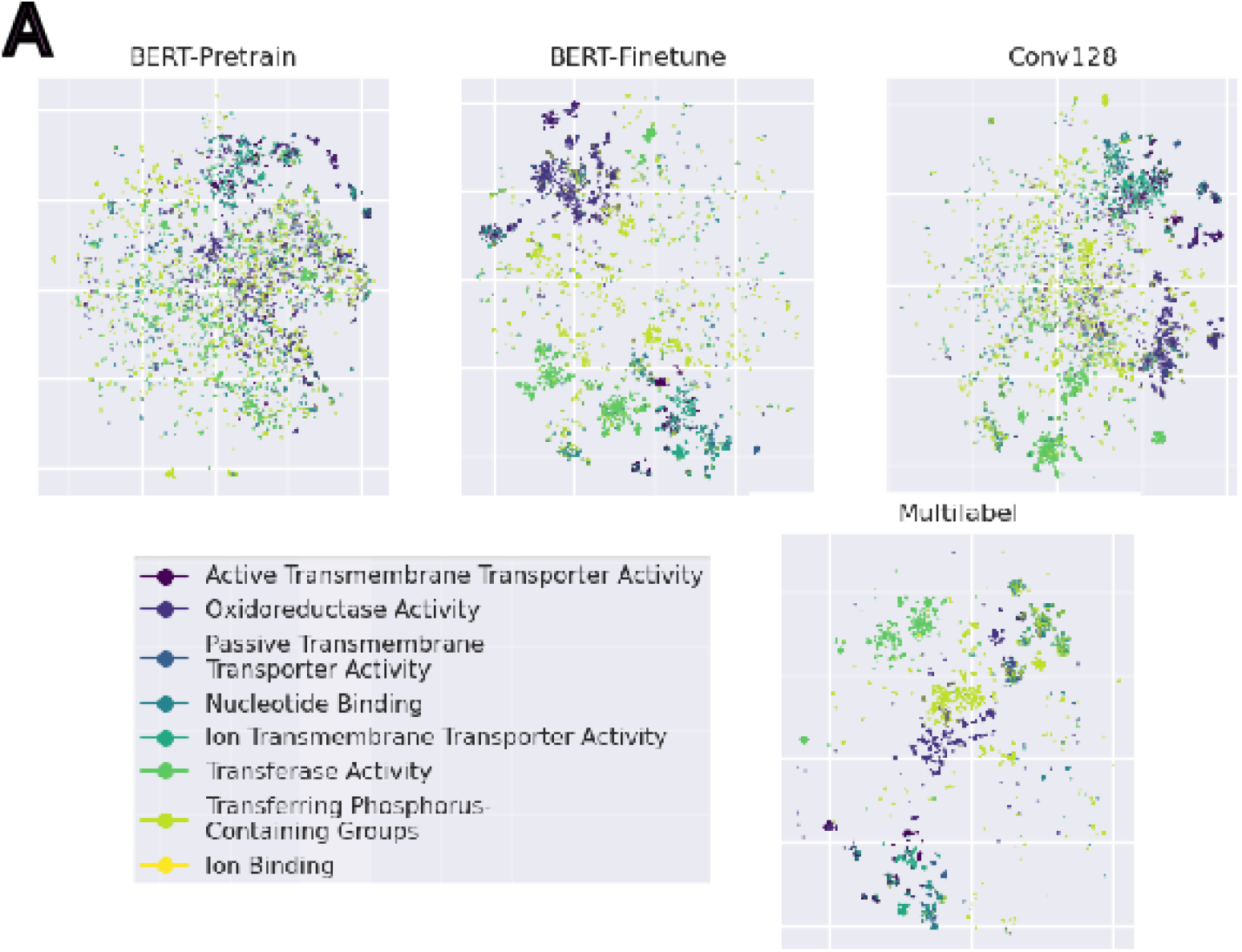
TSNE embedding of selected Gene Ontology classes. Classes are qualitatively selected to be high-level and relatively common, but reasonably distinct. Multilabel embedding is TSNE representation of actual protein labels, with no inference from sequence required, included for reference.

As protein annotation becomes an increasingly important task, effective, and actionable, annotation procedures become more valuable. We demonstrate that the improvement from fine-tuning is critical in this area, and that it has indirect benefits on the utility of protein representations, beyond improving direct classifier performance.

### Learning a Biologically Relevant Metric Space

Quantitative metrics on classification performance show that the fine-tuning produces a biologically meaningful space of embeddings. Nearby embeddings are likely to share functions, and simply propagating annotations between similar proteins outperforms deep learning alternatives.

TSNE visualizations of protein representations project them into two dimensions while attempting to maintain significant properties such as pairwise distance between neighbors (25, 26). TSNE scatterplots of ten significant GO classes indicate that GO classes are somewhat scattered among default pre-trained embeddings, but are indirectly clustered into functionally homogenous groups by fine-tuning classification loss. Qualitatively, fine-tuned embeddings look more organized than pre-trained or convolutional representations, or even TSNE representations built from label similarity.

### Rapid Queries of Protein Databases

An advantage of pre-computed protein embeddings over sequence is their high searchability in a database. Tools such as Faiss are pre-designed to provide rapid search of large vector databases, and allow for highly parallel, vectorized, searches of millions of protein embeddings in less than a second (27). When executing K-nearest neighbor queries, clustering, or other similarity based tasks, this can provide a significant computational advantage over sequence alignment.

### Benefit of Indirect Classification

It is not intuitively obvious, or typically true, that K-nearest neighbors approaches using embedding similarity would outperform machine learning classifiers trained on those same embeddings. Because the gradient descent optimization algorithm is explicitly focused on minimizing classification loss on logits calculated from the embedding, it would be expected that fine-tuning would optimize for this approach specifically. Our convolutional models do show this behavior, and are slightly more effective when classifying with logit predictions. However, embeddings from the fine-tuned language model consistently outperform logit predictions from the model.

One implication is that there is a significant qualitative difference between pre-trained language model embeddings, and typical embeddings from training on the annotation task exclusively. Additional information from pre-training may make embedding similarity a more useful heuristic, especially in low-data scenarios in the downstream task.

It remains difficult to explain the improved performance of embedding KNN classification over direct usage of models, but plots of precision and recall over GO classes give some hints. When we compare fine-tuned BERT with fine-tuned BERT embeddings, precision is similar across most classes, but recall is significantly improved by the usage of embeddings. It appears that for many classes, the deep learning classifier still almost completely fails to learn annotations for the majority of proteins, but that the similarity heuristic remains useful in these cases, significantly improving recall. Explicit inclusion of this heuristic in future models might close the gap, or surpass KNN methods. At the same time, the high performance of embedding or sequence similarity based heuristics suggests that current deep learning models don’t have a significant advantage over pattern matching based approaches, and are unlikely to extrapolate well to new functions or be significantly useful for design of novel functions.

## Materials and Methods

### GOBench Dataset

All experiments are run on a custom built GO (Gene Ontology) dataset from our previously developed gobench.org web application, with human-reviewed experimental codes (28) selected and otherwise default settings (29). The annotation dataset contains 112,000 unique, human-reviewed proteins, with an average of ten associated annotations out of 865 allowed possibilities.

### Language Model Fine-tuning for Gene Ontology Annotation

Finetuning is now a well-known strategy for boosting model performance in low-data regimes (6). We use the Rostlab ProtBERTBFD model, a transformer model pre-trained to predict masked amino acids in protein sequences from the BFD dataset (30), with roughly 250 million parameters.

ProtBERT uses a sequence of tokens representing protein residues as input, with a CLS token prepended to represent global information. BERT models are trained to predict masked sequences (31), so the output is a sequence of embeddings, with one embedding for each input token. However, our problem setting of GO classification requires global predictions for the entire protein, which requires modifying the ProtBERT structure.

Following fine-tuning convention, we average protein residue embeddings and add them to the model’s CLS embedding to get a single, 1024 dimensional, representation. We then linearly transform the embedding into 865 logit values, with one logit prediction for each GO class. This allows ProtBERT to function as a multi-class, multi-label classifier for GO terms, with some extra parameters from our final linear transform. During fine-tuning, ProtBERT is trained to minimize binary cross-entropy loss on GO class labels until convergence. We directly use the internal model embedding prior to the final classification layer as a protein representation.

### BLAST Annotation

We use BLAST as a strong baseline method for protein classification, based on the DiamondBlast method from DeepGOPlus (14, 32).

DiamondBLAST is a much faster variant of BLAST, which returns bitscores indicating similarity to each protein in a database, for each query protein in our testing dataset.

For a query protein *P*_*q*_ the score of a GO class *g* is an average of every associated database protein returned by DiamondBLAST, weighted by bitscore. Queries are run using default settings, which provides strong classifier performance in practice.

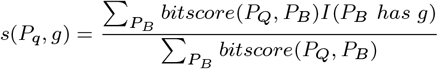

### Baseline Deep Learning Classifiers

We train deep learning classifiers from scratch as a point of comparison against fine-tuning large language models or BLAST. Deep convolutional models which aggregate large filters are known to be a strong alternative (14), so we select them as our predominant strategy, and use a reimplementation of the DeepGOPlus architecture to generate GO class predictions. As with our language models, we use internal model embeddings as protein representations.

### Baseline Optimization

Baseline machine learning models are trained until convergence, using the pytorch and the pytorch-lightning framework (33, 34).

DeepConv models require many important hyperparameters to be chosen, including learning rate, number of convolution kernels, maximum kernel size, and final embedding size, which may be very different for our particular GO dataset. We follow a Baysian optimization process for these four hyperparameters, and select our final values to maximize F1-score on our dataset after 80 trial training runs through Optuna (35).

### Multilabel KNN from Protein Embeddings

Protein sequences are converted to high-dimensional embeddings with one of our pre-trained, fine-tuned, or convolutional language models. Embedding size is 1024 for our pre-trained and tuned models, and 128 or 2048 dimensional for the convolutional models.

For KNN classification, all proteins in training and evaluation datasets are first embedded by some model. The training embeddings are treated as a reference database, with each protein embedding associated to a positive or negative assignment for each gene ontology class.

Metric distance is defined as the Euclidean distance between vectors. For each evaluation protein, we search for the K training embeddings with the smallest distance. Then for each Gene Ontology class, we estimate the class probability by the percentage of nearest-K training embeddings positively annotated with this class. For our problem setting, we find that performance is relatively robust to the specific value of *K*, and adopt *K* = 10 for our evaluation. We also find that reweighting neighbors by their distance makes little overall difference to final performance.

## ACKNOWLEDGMENTS

We thank Salmonn Talebi, Ehsaneddin Asgari, and other members of the Molecular Cell Biomechanics Laboratory for the suggestions and comments they provided throughout this research.

